# Pancreatic adenocarcinoma human organoids share structural and genetic features with primary tumors

**DOI:** 10.1101/338897

**Authors:** Isabel Romero Calvo, Christopher Weber, Mohana Ray, Miguel Brown, Kori Kirby, Rajib K. Nandi, Tiha M. Long, Samantha M. Sparrow, Andrey Ugolkov, Wenan Qiang, Yilin Zhang, Tonya Brunetti, Hedy Kindler, Jeremy P. Segal, Andrey Rzhetsky, Andrew P. Mazar, Mary M. Buschmann, Ralph Weichselbaum, Kevin Roggin, Kevin P. White

**Author notes:** CORRESPONDING AUTHOR: Kevin P. White, Ph.D, Chicago Pancreatic Cancer Initiative, Institute for Genomics & Systems Biology, The University of Chicago, 900 E. 57^th^ Street, Chicago, IL and Tempus Labs 600 West Chicago Avenue, Chicago, IL. or. **CONFLICT OF INTEREST** Kevin White, Andrey Ugolkov and Yilin Zhang are employees of Tempus labs; Andrew Mazar is the founder of Tactic Pharma, Pamdeca, Actuate Therapeutics, Lung Therapeutics, Inc, and Monopar Therapeutics; All other authors have nothing to disclose.

## Abstract

Patient-derived pancreatic ductal adenocarcinoma (PDAC) organoid systems show great promise for understanding the biological underpinnings of disease and advancing therapeutic precision medicine. Despite the increased use of organoids, the fidelity of molecular features, genetic heterogeneity, and drug response to the tumor of origin remain important unanswered questions limiting their utility. To address this gap in knowledge, we created primary tumor- and PDX-derived organoids, and 2D cultures for in-depth genomic and histopathological comparisons to the primary tumor. Histopathological features and PDAC representative protein markers showed strong concordance. DNA and RNA sequencing of single organoids revealed patient-specific genomic and transcriptomic consistency. Single-cell RNAseq demonstrated that organoids are primarily a clonal population. In drug response assays, organoids displayed patient-specific sensitivities. Additionally, we examined the *in vivo* PDX response to FOLFIRINOX and Gemcitabine/Abraxane treatments, which was recapitulated *in vitro* by organoids. The patient-specific molecular and histopathological fidelity of organoids indicate that they can be used to understand the etiology of the patient’s tumor and the differential response to therapies and suggests utility for predicting drug responses.

## INTRODUCTION

In the United States alone, pancreatic cancer is the 3rd leading cause of cancer death, accounting for over 50,000 cases annually, with a survival rate of less than 10% (1). The most common form of pancreatic cancer, pancreatic ductal adenocarcinoma (PDAC), is often diagnosed at a late stage, limiting effective therapeutic interventions. Only 10–15% of patients are eligible for surgery, the only potentially curative option (2). Although adjuvant chemotherapy has shown incremental improvements in resected patients, the vast majority recurs and succumbs to metastatic disease with a five year survival of only 1-2%. This poor survival underscores the need for novel tools to rapidly and precisely match patients with effective therapies.

The limited number of available preclinical PDAC models has been a major hurdle for discovery and translational research. Development of *in vitro* cell culture models, as well as transgenic and patient derived xenograft (PDX) mouse models, enables investigation of biological mechanisms and new treatments. However, traditional two-dimensional (2D) immortalized monolayer cell lines are limited, in that they do not reflect the variability or the structure of PDAC tumors. Primary cells cultured as monolayers do not reflect the heterogeneity of the primary tumor due to selection in culture, lack of stromal-stromal-cell communication, tissue-specific architecture and mechanical cues. Combined, these impact impact gene expression, which is key for modeling therapeutic responses (3-5). Genetically-engineered mice, such as *LSL-Kras^G12D/+^*; *LSL-Trp53^R172H/+^*; and *Pdx-1-Cre* are useful in that they are models of carcinogenesis within the pancreas, however they only represent very specific genetic mutations and fail to recapitulate the variability that is characteristic of PDAC (6-8). Furthermore, PDX models have complex architecture and utilize heterogeneous patient tissue, but the clinical applicability is challenging due to prohibitive time requirements (up to 6 months to establish tumor growth), costs associated with *in vivo* systems, the large sample size required (~100 mm^3^), and the influence of infiltrating murine stromal cells on the tumor (9, 10). The latter factor may influence the recent observations that the more times a PDX tumor is passaged through mice, the more transcriptionally “mouse-like” it becomes (11). Due to the challenges related to existing systems, improved models of PDAC are essential.

Recently developed three-dimensional (3D) organoids may provide dramatic benefits compared to 2D cell culture and PDX models for their use in personalized medicine and drug discovery. Organoid models are rapidly and inexpensively developed from primary or metastatic tumor tissue for expansion and molecular profiling. Organoids also recapitulate biological features of a 3D environment, as demonstrated in pancreatic, colorectal, prostate and mammary cancers (12) (13) (14) (15). Importantly, PDAC organoids can be grown with a high rate of success from very small amounts of tissue collected from diagnostic fine-needle biopsies (16, 17), core needle biopsies, and in excisional intraoperative biopsies and these tissues are more likely to be passaged over time when grown in 3D versus 2D cultures (18) (19) (16) (17).

Some barriers to the use of organoids in personalized therapies include lack of in-depth assessment of the histopathology, genetic stability, and molecular profiling of organoid models. Comparisons are needed between organoids and primary tumors of origin, as well as among individual organoids derived from the same patient. Before organoids can be routinely employed as model systems more characterization of cellular and molecular properties are required. Specifically, insufficient morphological characterization of organoids has been reported and comparisons are needed between organoids and primary tumors of origin, as well as among individual organoids derived from the same patient. Therefore, we performed histopathological profiling on organoid and PDX models for comparison to their corresponding primary tumors. In addition, we performed a thorough genomic characterization of organoid cultures by deep sequencing to determine if the models represent the genomic constitution of the primary tumor, and if they retain their genomic characteristics over time. We performed *ex vivo* drug testing to improve our understanding of the degree to which organoids model the therapeutic response of the corresponding tumor of origin. We have defined the histopathology, genetic heterogeneity and therapeutic sensitivity profiles of organoid models derived from PDAC patients as an initial effort to provide thorough pathological and genetic comparison between PDAC organoid models and the tumors from which they are derived.

## MATERIAL AND METHODS

### Human specimens

Between 2014 and 2017, 10 tumor samples were collected from patients with pancreatic cancer adenocarcinoma. The clinical information from those patients was collected (Supplementary Table 1). All patients provided informed consent and the study was conducted under IRB12-1108 and IRB13-1149. Clinical data was obtained from the electronic medical record (Epic). Pancreatic ductal adenocarcinoma and adjacent uninvolved pancreas were obtained from patients undergoing pancreatic resections at The University of Chicago Medicine (UCM) facilities. Samples were confirmed to be tumor or benign based on pathological assessment. Normal pancreas tissue was obtained from patients who underwent resection for benign lesions and demonstrated no features of pancreatitis.

### Patient-Derived-Xenograft Mice (PDX)

The research protocol was performed under IS00000556 and IS00000424 Institutional Animal Care and approved by Northwestern University. Human tumor samples were obtained from UCM pancreatic cancer patients and de-identified. In brief, freshly resected human tumor samples (0.2 g) were sent to Northwestern University and transplanted subcutaneously into non-obese diabetic/severe combined immunodeficient (NOD-SCID) gamma (NSG) mice (Jackson Laboratory). Tumor volumes (length*width^2^)/2) were measured weekly. Default endpoints for any animal was loss of 15% body weight compared to pre-tumor weight, tumor ulceration, clinical or behavioral signs including lack of responsiveness, inactivity, hunched posture and unresolved skin ulcers. When PDX tumor reached 1.5 cm in its largest dimension, the mouse was euthanized and freshly resected 2mm × 2mm tumor pieces (avoiding necrotic regions) were retransplanted to NSG mice for *in vivo* studies. A piece of subcutaneous PDX tumor was fixed in 10% formalin and processed to paraffin-embedding. Paraffin sections (5 μm) were stained with hematoxylin and eosin (H&E) for histopathological evaluation.

For initial treatment studies, mice bearing subcutaneous pancreatic PDX tumors from patient 1 were staged to approximately 200 mm^3^ prior to initiation of treatments and randomized to two treatment groups: control (vehicle) and FOLFIRINOX (4 mg/kg oxaliplatin, Zydus Hospira, 61703-363-22; 50 mg/kg leucovorin, Teva, 0703514501; 25 mg/kg irinotecan, Pfizer, 0009752903; 50 mg/kg 5-FU, Thomas Scientific, F6627; n=3, 2 tumors per mouse). Vehicle or drugs were injected intraperitoneally once a week for 3 weeks.

For the next set of studies, mice bearing subcutaneous pancreatic PDX tumors from patient 1 were staged to approximately 150 mm^3^ prior to initiation of treatments and randomized to 3 treatment groups: control (vehicle), gemcitabine (100 mg/kg, Sigma, G6423) and gemcitabine (100 mg/kg) + abraxane (30 mg/kg, Celgene, 6881713450); n=3, 2 tumors per mouse. Vehicle or drugs were injected intraperitoneally once a week for 4 weeks.

### Isolation and culture of human pancreatic cancer organoids and 2D cells

To establish 2D cells and PDX-derived organoid cultures, a piece of subcutaneous PDX tumors was extracted, minced and digested with collagenase type XI (0.125 mg/ml, Sigma, C9407) and dispase (0.125 mg/ml Gibco, 17105041), in DMEN (Gibco, 11995065) and incubated from 0.5-1 h at 37 °C with gentle shaking. Cells were spun and dissociated with TrypLE Express (Fisher Scientific, 12605-010) and DNase I, 10 μg/ml (DN25, Sigma), 10 min at 37 °C, and washed with DMEN. For culturing 2D cells, an aliquot (1/10) of the dissociated suspension was seeded into a petri dish with complete 2D media (DMEN-F12 Advanced, GIBCO, 12634-010; HEPES buffer, Invitrogen, 15630-080; penicillin/streptomycin, Thermo Fisher, 15140-122; L-GlutaMax, Invitrogen, 35050-061; FBS 5% Life Technologies, 13028-014; EGF 10 ng/ml, GIBCO, PMG8043; Bovine Pituitary Extract ~57.6 μg/ml, hydrocortisone 2μg/ml, Sigma, H0888 and insulin human recombinant 0.56 μg/ml, Life Technologies, 12585-014). For culturing organoids, dissociated cells were washed and embedded in growth-factor-reduced (GFR) Matrigel (Corning, 356231) and cultured in complete media (Intesticult [Stemcell Technologies, 6005], A83-01 [0.5 μM, Sigma, SML0788], fibroblast growth factor 10 [FGF10, 100 ng/ml, Gibco, PHG024)], Gastrin I [10 nM, Sigma, 17105-041], N-acetyl-L-cysteine [10 mM, Sigma, A9165], Nicotinamide [10 mM, Sigma, N0636], B27 supplement [1x, Gibco, 17504-044], Primocin [1 mg/ml, InvivoGen, ant-pm-1] and Y-27632 [10.5 μM Tocris, 1254]. To establish tumor-derived 2D cell lines and organoids, resected primary tumor samples from pancreatic resection surgeries were utilized as above. Organoids were passaged via mechanical dissociation with TrypLE Express (Fisher Scientific, 12605-010) and passage was preformed weekly with a 1:2 ratio.

### Quantification of H&E architecture

A gastrointestinal pathologist scored architecture of tumor and organoids from H&E stained slides according to the three different patterns (**Figure 1b**). The patterns observed were i. Simple (score=1): tumor epithelium composed of a single layer of epithelial cells, but often with some loss of nuclear polarity, ii. Papillary (score=2): tumor cell composed of focal stratification into multiple layers with occasional luminal projections, iii. Solid/cribriform (score=3): tumor cells grow in a syncytium, within occasional small gland structures. An average aggregate score was calculated from the total number of fields and the percent of fields containing each of the three patterns, according to the following equation: Histology score = [Fields of pattern 1 + Fields of pattern 2 + Fields of pattern 3]/total fields.

**Figure 1.**
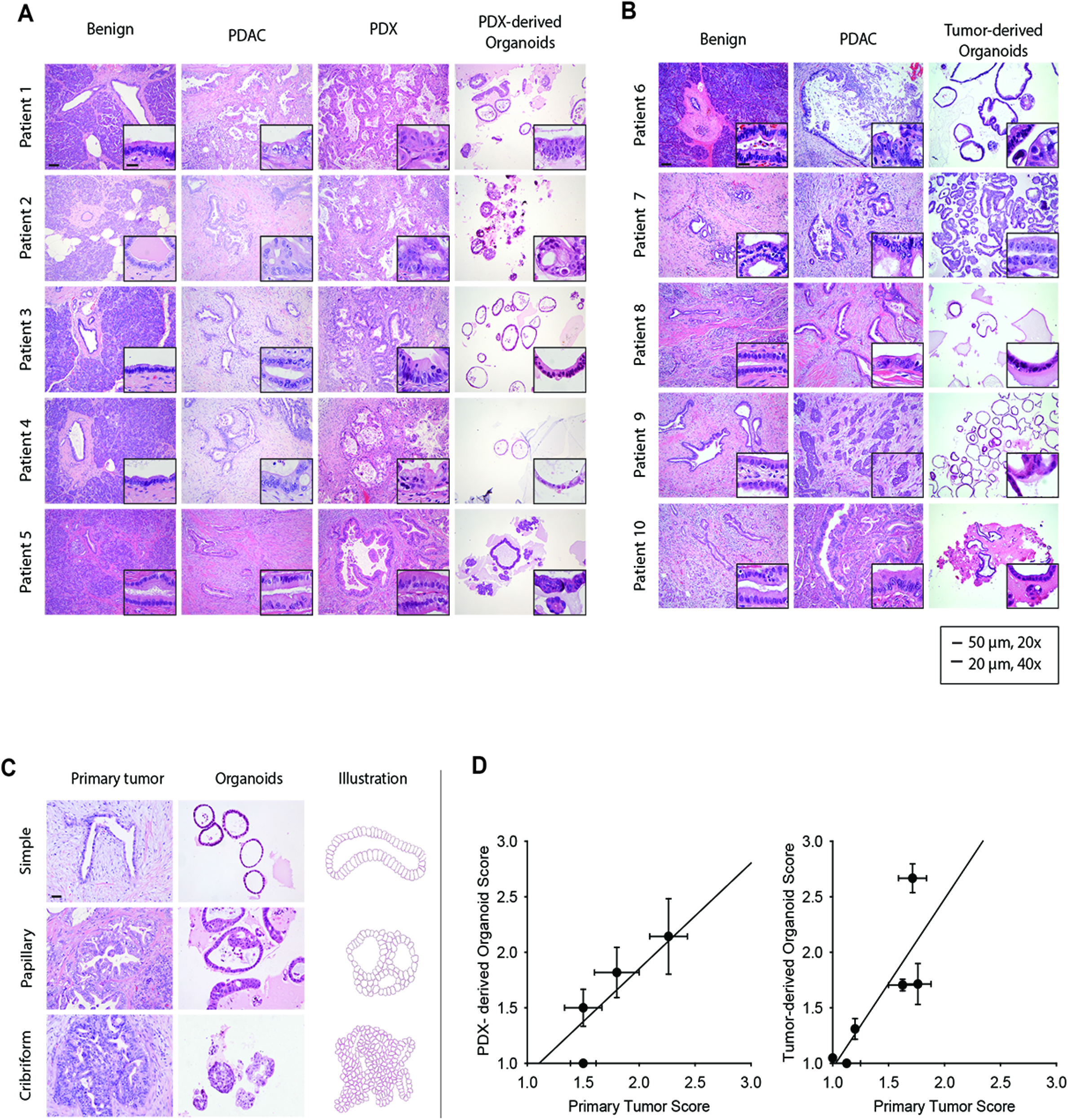
Patient and PDX-derived organoids share morphological features with the primary tumor. (**A**) Hematoxylin and eosin (H&E) stained slides show glandular architecture of primary tumors, PDX, and PDX-derived organoid models, patients 1-5 (n=5), and (**B**) patient-derived organoids, patients 6-10 (n=5). (**C**) Primary tumors and organoid morphologies were classified as *Simple, Papillary*, or *Cribiform*. Representative images are shown (left) and a diagram (right). (**D**) H&E stained slides were utilized to quantify the correlation between primary tumors and PDX-derived organoid models, patients 1-5 (n=5), R^2^=0.67, and tumor-derived organoids, patient 6-11, R^2^=0.77 (E). Representative images selected and scored by gastrointestinal pathologist. Imaging at 20X (bar 50μm), inset at 40X (bar, 20μm).

### Immunohistochemistry staining and quantification

Human samples, PDX and organoids were fixed with 10% formalin, paraffin embedded and sectioned (5μm). The following antibodies were used: CK7 (Agilent/DAKO, M7018, mouse monoclonal, clone OV-TL 12/30), CK19 (Agilent/DALO, M0888, mouse monoclonal, clone RCK108), CK20 (abcam, ab76126, rabbit IgG), CEA (abcam, ab4451, mouse monoclonal, clone 26/3/13), Claudin-4 (abcam, ab53156, rabbit IgG.), CA19-9 (Thermo Fisher, MA5-13275, mouse monoclonal, clone 121 SLE) and P53 (Calbiochem, OP-43-100UG, mouse monoclonal, clone DO-1). Slides were scanned (ScanScope XT, 20x). Quantification was performed using Imagescope (Aperio eSlide Manager Sofware, Leica). Staining intensity was scored 0 to 3 using the positive pixel count algorithm.

### DNA extraction and library preparation

#### Primary and PDX tumors

A GI pathologist scored H&E sections of the tumor and circled areas of high tumor cellularity. DNA was extracted from the same area of an adjacent normal section using the QIAamp DNA Micro kit (56304), quantified and processed into indexed libraries using the Illumina TruSeq kit (FC-121-2001) following the manufacturer’s protocol.

#### 2D cell line and organoids

For 2D cell lines, DNA was extracted from approximately 3 million cells using the DNeasy Blood and Tissue kit (Qiagen, 69504) following manufacturer’s protocol. DNA was extracted from a single organoid for analysis by DNA-seq. The Matrigel in one well of a 24-well tissue culture plate was depolymerized with 4°C PBS. Under 4X brightfield magnification, a single organoid was isolated in 3-4 uL of liquid Matrigel, and DNA was extracted and amplified as recommended by the manufacturer. The GenomePlex Single Cell Whole Genome Amplification Kit (Sigma Aldrich, WGA4-10RXN) and the Repli-G Single Cell kit (Qiagen, 150343) were utilized. Following amplification, the DNA Clean and Concentrator (Zymo Research, D4013) kit was utilized and libraries were generated using KAPA LTP Library Preparation kit (KAPA Biosystems, KR0453).

#### Targeted capture and sequencing

The indexed libraries were pooled, and targeted capture of the exons of 1,213 known cancer genes was carried out (20). In brief, this approach utilizes a tiered assay system in which highly clinically relevant genes (tier 1, n=316) are sequenced approximately 2.18-fold deeper than the remaining (tier 2) genes. The enriched libraries were sequenced (101 bp paired-end reads) using Illumina HiSeq and FastQ files were generated using BCL2FastQ1.8.4 (Illumina Inc.)

### DNAseq analysis pipeline

#### Quality control and read alignment

FastQ files were trimmed using Cutadapt v1.9.1 (http://cutadapt.readthedocs.io/en/stable/guide.html) to remove low quality bases and adapter sequences. The reads were aligned against build 19 of the human genome reference (hg19) using BWA-MEM v0.7.8 (http://bio-bwa.sourceforge.net). For PDX samples, reads arising from mouse stroma were filtered out by mapping against a custom-built genome of the mouse strain used for PDX generation, and removing all perfectly aligned reads, prior to human reference alignment. PCR duplicates were removed while sorting on-the-fly using novosort v1.03.9 (Novacraft Technologies Sdn Bhd, http://www.novocraft.com/products/novosort/). Bedtools v2.22.1 (http://bedtools.readthedocs.io/en/latest/) was used to ascertain coverage at tier 1/tier 2 loci. A threshold of 300X mean coverage at tier1 genes was used for tumor, normal, PDX and 2D cell line samples, while organoid samples were allowed more leniency in tier 1 coverage (median coverage 226x).

#### Variant calling

Single nucleotide variants (SNVs) were called using MuTect v1.1.7 (http://archive.broadinstitute.org/cancer/cga/mutect) while insertions and deletions (indels) were called using scalpel-discovery 0.5.3 http://scalpel.sourceforge.net/). Indel calls not annotated as “PASS” or “KEPT” were removed. For both SNVs and indels, only calls within genomic regions targeted by the capture panel were retained for subsequent analyses. All variants were annotated using ENSEMBL’s Variant Effect Predictor version 84 (http://www.ensembl.org/info/docs/tools/vep/script/vep_download.html) and only variants within coding regions or disrupted splice sites were included in analyses. Calls with a variant allele frequency (VAF) < 5% for the patient tumor and normal samples, position coverage < 30, or an allele frequency >= 0.01 in ExAC (http://exac.broadinstitute.org/) were removed. Exceptions were made in cases where either a low frequency variant was called in the models and present at less than 5% VAF in the tumor, and when a variant was called in the tumor but just missed the cutoff in the models. To further improve the quality of indel calls, low complexity genomic regions were identified using Dustmasker (https://www.ncbi.nlm.nih.gov/IEB/ToolBox/CPP_DOC/lxr/source/src/app/dustmasker/) a pseudo panel of normal samples was constructed by grouping putative indel calls that failed Scalpel filters due to ‘HighVafNormal’ or ‘HighAltCountNormal’. All indels that failed in two or more samples from unique patients were removed.

### RNA Extraction, library preparation and sequencing

#### Primary and PDX tumors

A GI pathologist circled areas of high tumor cellularity on H&E sections of the tumor and PDX. RNA was extracted from the same area of an adjacent section using RNeasy FFPE kit (Qiagen, 73504) after treating with Deparaffinization Solution (Qiagen, 19093).

#### 2D cell line and organoids

The RNeasy Mini kit (Qiagen, 74104) was used for extracting RNA from both 2D cell lines and organoids. Approximately 3 million cells and 1 million cells were used as starting material for 2D and organoids, respectively.

#### RNA library preparation and sequencing

Exome capture was carried out using the RNA access kit (Illumina, RS-301-2001) for the majority of samples, with the exception of 2D cell lines from patients 1-3 which had total RNAseq performed with ribosome depletion (Illumina, MRZH11124). Libraries were pooled and sequenced on HiSeq or NextSeq 500 instruments, to generate paired-end 100 bp reads.

### RNASeq analysis pipeline

Sample specimens from tumors, normals and models from patients 1-4 and 7 were quantified using RNAseq. The cell lines in the multi-patient comparisons were sequenced using totalRNAseq with ribosome depletion (Illumina, MRZH11124), while FFPE derived tissue samples were sequenced using the RNAaccess capture kit from Illumina (RS-301-2001). All samples were quality checked and aligned using the following general outline: Reads from fastq files were adapter and quality-trimmed with a minimum phred score of 20 on 3’ ends using cutadapt v 1.15 (21) and read quality was evaluated further using FastQC (https://www.bioinformatics.babraham.ac.uk/projects/fastqc/Andrews, 2014). Reads were aligned using STAR v 2.4.2a (https://github.com/alexdobin/STARcite Dobin et al., 2013). Alignment quality was assessed using Picard v2.16 CollectRnaSeqMetrics for read counts, percent unique mapping reads, percent intergenic reads, and percent ribosomal ref ads to assess depth and capture quality. Samples were then quantified using eXpress (22). A minimum of 20 million reads per sample was required for quantification and subsequent differential expression analysis. TPM values were obtained from eXpress output and collapsed on gene name and limited to protein coding and lincRNA for multi-patient quantification comparison scatterplots. For differential expression, effective counts were used from eXpress output and similarly collapsed on gene name and limited to protein coding and lincRNA for patient 1 comparisons.

The R princomp and DESeq2 (23) libraries were used for differential gene expression analysis using the effective counts output from eXpress. PCA combined with linear regression was used to determine which covariates should be included in the final regression model for differential expression. In the set of samples used for multi-patient comparison, no significant covariates were identified besides the independent variable of interest. However, for the indepth analysis of patient 1, there was a detectable batch effect influencing principal component 1 (adjusted r squared 0.758, p-value 0.00028), and was included as a covariate in the general linear model using DESeq2. Genes were considered differentially expressed if they had an adjusted p-value < 0.05 and a log2 fold change greater than 1 or less than −1. A representative subset of the top differentially expressed reads by adjusted p-value was generated by selecting the top 200 differentially expressed genes, defined as genes with the lowest adjusted p-value with a minimum mean coverage of 105.6608 from each biotype (tumor, PDX, 2D cell line, organoid). The union of these top 200 genes, from each biotype was generated for a total of 518 unique genes that were considered in downstream analysis. The log2 fold change values from the genes in the union were extracted and any gene in the union that had an adjusted p-value > 0.05 in a particular biotype was not considered differentially expressed and therefore the log fold change was not taken into consideration for that biotype. For the multi-patient comparison, since sequencing was done over two lanes each for each patient, each lane was treated as a replicate to increase accuracy and power. Multiple runs were used in the in-depth analysis except for the PDX. Data from normal patient tissues in the study were combined and used as controls for all differential expression analyses.

### Drop-Seq single cell RNA-Seq

The single cell RNA-seq dataset is comprised of 15 separate drop-seq runs. For each run, a total of 2000 mature organoids were pooled, typsinized with TrypLE in presence of 10 μM Y-27632, strained into a single cell suspension and counted. The final concentration of the cell suspension varied between 100-120 cells/uL depending on the flow rate at which monodisperse droplets were formed. An aliquot from the cell suspension was stained with trypan blue to ensure high cell viability (>90%) at the time of establishing the cell preparation protocol. Organoids were harvested for drop-seq analysis at same growth size to ensure uniformity from run to run.

Nanoliter-sized droplets containing single cells and barcoded beads were prepared as described in Macosko et al. (24). Briefly, droplets generated by co-flowing 1 mL each of cell and bead solutions through a microfluidic device were collected. The droplets were broken and the beads were collected and reverse-transcribed. The cDNA obtained was then PCR amplified and quantified. Finally, the cDNA was fragmented and amplified using primers that allow amplification of only the 3’ ends, processed into RNA-seq libraries and sequenced on the Illumina NextSeq 500 sequencer. The sequencing data was analyzed using the pipeline created by Macosko et. al. (24) to generate digital gene expression (DGE) matrices. Only cells with more than 250 genes were retained for subsequent clustering.

We developed a novel clustering approach based on a class of probabilistic generative models called topic models. In brief, the DGE is first normalized, scaled and log transformed. The contribution of undesirable sources of variation such as cell cycle phase, batch effect and mitochondrial gene expression is assessed by calculating scores for each of these factors and regressed out of the data (25). Next, a training set comprised of cells that express at least 900 genes is created and highly variable genes are identified (26). A topic model is then generated from the training set using the highly variable genes as the vocabulary (27). Then, a topic distribution is calculated for the cells not included in the training set by comparing their gene expression to that of the training set. Statistical significance of topics is calculated based on KL-divergence and insignificant topics are eliminated. Finally, cells are assigned to the topic with the highest probability, and clusters with fewer than 50 cells are removed. Pairwise marker genes for each cluster are identified using genes that are expressed in least 10% of the cells in both clusters and have average log fold change >1 between clusters. Clusters that do not have at least 10 unique marker genes are merged (28).

### Organoid treatment experiments

#### Real time viability

Relative viability of organoids was determined by measuring Annexin V fluorescence in real-time with Incucyte (Essen Bioscience). Organoids were seeded (3000 cells/well) into a U-bottom ULA 96 well-plate (Costar, 7007). Cells were incubated with Annexin V Green Reagent (Essen Bioscience, 4642) and treated with Gemcitabine Hydrochloride (G6423, Sigma) at 3nM, 10nM. and 30nM. Real-time fluorescence was meassured with Incucyte during 72 hours.

#### Measurement of cell viability (MTA assay)

Relative number of viable cancer cells was determined by measuring the optical density using CellTiter 96 Aqueous One Solution Cell Proliferation Assay kit [3-(4,5-dimethylthiazol-2-yl)-5-(3-carboxymethoxyphenyl)-2-(4-sulfophenyl)-2H-tetrazolium, inner salt; MTS] according to the manufacturer’s instructions (Promega, G3580, Madison, WI). To resemble clinical pharmacokinetics of SOC drugs, tumor organoids were treated with vehicle, gemcitabine, gemcitabine and paclitaxel, and FOLFIRINOX (oxaliplatin, leucovorin, irinotecan, 5-fluorouracil) for 4 hours at indicated concentrations. Post-treatment, test compounds were replaced with fresh compound-free complete culture media. Organoids were grown another 72 hours and relative cell growth and death (as compared to optical density signal before the start of the treatment) was measured by MTS cell viability assay after 72 hours of growth.

### Statistical analysis, differential expression

For differentially expressed genes, R PCA package princomp was used to check for batch effects. In the set of samples used for multi-patient comparison, batch effects were negligible and therefore were not regressed out. However, for the in-depth analysis of patient 1, there was a detectable batch effect influencing principal component 1 (adjusted r squared 0.758), so it was included in the batch model for DESeq2. Differential expression with p values and adjusted p values were calculated using default methods of negative binomial global linear model and Wald statistics. For the multi-patient comparison, since sequencing was done over two lanes each for each patient, each lane was treated as a replicate to increase statistical significance; for the indepth comparison, for all but PDX there were multiple runs (2-4). In all cases, normal tissues from all patients in the study were used as the control for differential expression. Differentially-expressed genes were analyzed by Ingenuity Pathway Analysis to predict cancer related, The Benjamini-Hochberg method was utilized and an FDR cutoff of q<0.05 was applied. 2way ANOVA Multiple comparisons were performed in all *in vivo/in vitro* studies.

### Data availability

The genomic data from this publication has been deposited at www.ncbi.nlm.nih.gov. The submission ID is SUB4032998. The BioProject ID is PRJNA471134. Private login credentials are pending with the NIH.

## RESULTS

### Organoids from both primary PDAC and PDX share morphological features with the primary tumor

In order to generate clinically relevant PDAC models, benign and tumor tissue were collected at the time of surgical resection (**Table S1**). PDAC organoids were grown from both PDX and primary tissue (15). PDAC tumors from five patients were used to establish PDX tumors, followed by the growth of PDX-derived organoids from those xenografts. In addition, PDAC tumors from another five patients were utilized to grow primary tumor-derived organoids directly (without PDX) (**Figure S1**).

To validate tumor pathology in PDX and organoid models, hematoxylin and eosin stain (H&E) histological analysis of uninvolved pancreas, primary tumor, PDX and organoid models was performed. A gastrointestinal pathologist compared the architecture and cell morphology in both PDX and organoid models to the primary tumors (**Figure 1A**). Benign ducts revealed simple columnar epithelium without significant cytological atypia, and areas of PDAC epithelium showed various degrees of architectural disorder and cytological atypia. Specifically, tumors from patients 3 and 5 (**Figure 1A**) showed well differentiated tumor glands with small basally oriented nuclei and little or no cellular stratification. Tumor glands from patient 4 were slightly more atypical and demonstrated regions of stratification. Glands from patients 1 and 2 showed the greatest extent of cell atypia and cell stratification. We divided the architectural patterns of the organoids into three morphological classifications: simple, papillary, and solid/cribriform (**Figure 1C**). These architectures were quantified, and a high degree of correlation was observed between the morphological structure of the primary tumors and the corresponding organoids (R squared=0.67) (**Figure 1D, left**). The architecture of the PDX models was more complex and atypical compared to the organoid models, in part due to the presence of mouse stroma within the tumor. Nevertheless, organoids derived from the PDX models recapitulated the microscopic features of the primary tumors.

To determine if organoids derived directly from the primary tumor also maintain tissue architecture similar to PDX-derived organoids, we grew organoids directly from the primary tumor (patients 6-10) and performed histological analysis of tissue derived from these five patients. We observed a similar glandular architecture between primary tumor and organoid models (**Figure 1B**). Like those derived from PDX passaged tumors, organoids derived directly from primary tumors maintained the morphological structure of their corresponding primary tumor. Specifically, the well-differentiated tumor glands seen in the organoids from patients 8 and 10 resembled the relatively simple glandular architecture observed in the primary tumors from these patients. Likewise, for patients 6, 7, and 9, the organoids demonstrated more cellular stratification akin to the primary tumors of these same patients. Quantitative analysis of glandular architecture demonstrated a high degree of morphological correlation between primary PDAC and organoids (**Figure 1D, right**) (R squared=0.77).

### Organoids and patient-derived tumor xenografts share protein expression features with corresponding primary tumors

To define the relevant protein expression profiles of the PDX and organoid models and to compare these protein expression patterns with both benign pancreatic and tumor tissue, we used a panel of immunohistochemical (IHC) markers. Since the majority of benign tissue obtained from the tumor resections demonstrated evidence of chronic pancreatitis, we also stained normal pancreatic samples selected from patients without PDAC and with no evidence of pancreatitis. Cytokeratin markers 7 and 20 (CK7 and CK20) were selected to assess for pancreatic epithelial differentiation, and cellular tumor antigen p53 (p53), Claudin-4 and Cancer Antigen 19-9 (CA19-9) were selected due to their frequent expression in neoplastic pancreatic ductal epithelium (29-32) (**Figure 2A, 2B**).

**Figure 2.**
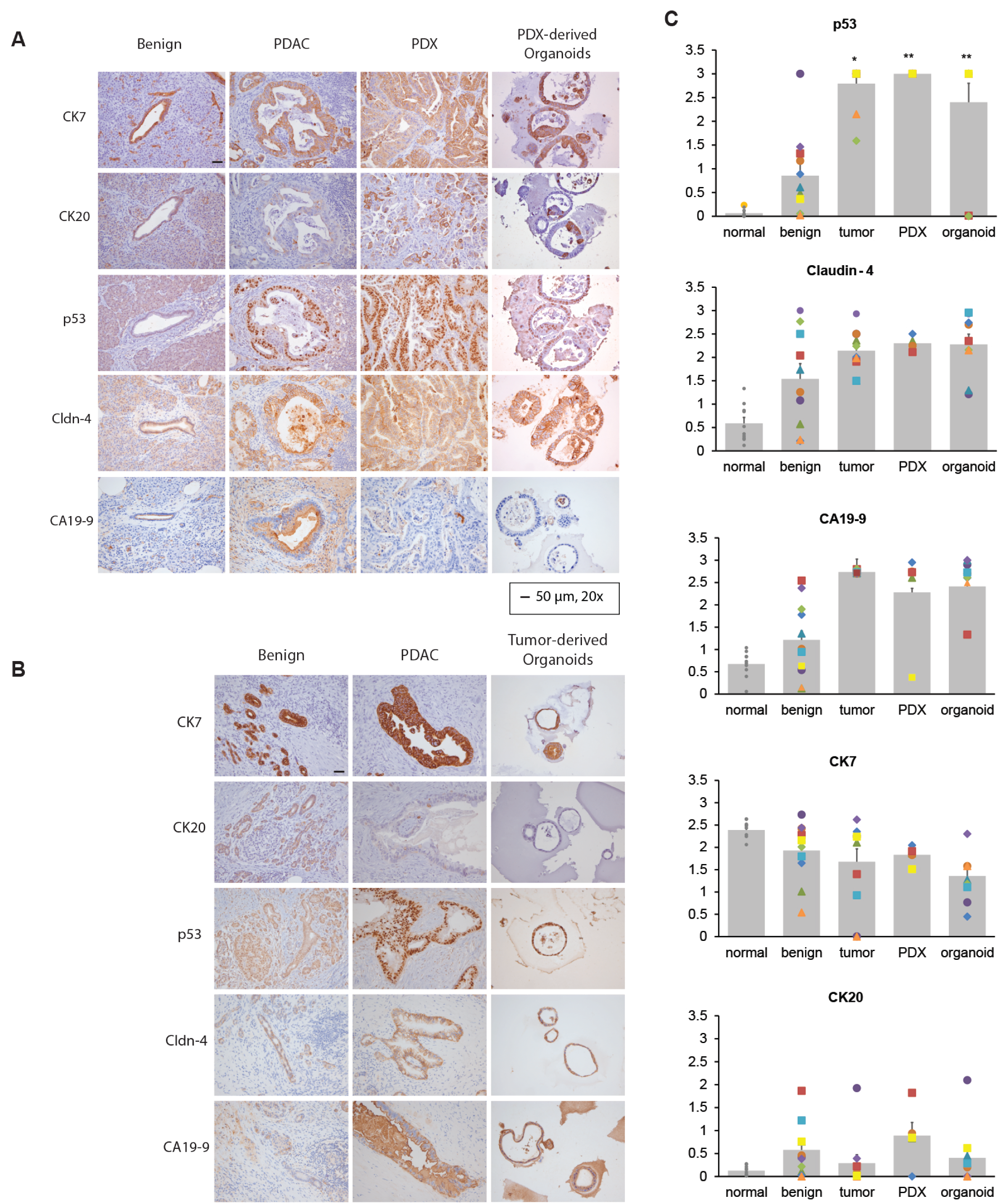
Patient and PDX-derived organoids share protein expression features with the primary tumor. (**A**). Representative IHC staining patterns from patients 1-6 of CK7, CK20, p53, CEA, Claudin-4, and CA19-9 of primary PDAC, PDX and PDX-derived organoid models (patients 1-6), and (**B**) patient-derived organoids (patients 6-10). Results are representative of 5 independent fields. Imaging at20X imaging (bar 50mm). (**C**) CK7, CK20, p53, Claudin-4 and CA19-9 protein levels from normal and benign pancreas, primary tumor, PDX, and organoids were quantified. Average expression (gray bar) was calculated by quantifying pixel intensity of 5 different fields per patient ŕn=10, individual patients reflected bv independent shapes and colors), *p<.05 versus benign, ** p<.05 versus adjacent normal.

To quantify the extent that protein expression in the organoid and PDX models was recapitulated by the staining observed in the primary tumor, pixel count and intensity were measured for all cases. Cytokeratin proteins, normally found in the intracytoplasmic cytoskeleton of epithelial tissue, change their protein profile expression in PDAC. CK7 and CK19 are expressed in 90-100% of PDAC (33, 34). In contrast, CK20 is found in less than 20% of PDAC cases (35). We found that the protein expression profiles of these cytokeratins is maintained in the PDX and organoid models (**Figure 2, Figure S2**). We also examined nuclear labeling of the mutated tumor suppressor p53, as aberrant expression occurs in at least 70% of PDAC (31, 36). Consistent with these previous observations, we found that normal pancreatic epithelium showed low levels of p53 expression while p53 is increased in primary tumors and corresponding PDX and organoid models (**Figure 2**). We also stained for Claudin-4, which is often overexpressed in PDAC (37) (38), and we observed that its pattern of overexpression is elevated in both PDX and organoids models (**Figure 2**). Next, we examined tumor epithelial cells for the expression of the sialyl Lewis motif that corresponds to CA19-9 (39). We found that all of the primary tumors studied expressed CA19-9, and that both the PDX and organoid models stained positive for CA19-9 as well. In normal pancreatic epithelium, we saw that CA19-9 was restricted to the luminal surface. However, in tumor cells, the antigen was localized in the cytoplasm, basolateral membrane, and occasionally at the apical cell membrane. In addition, the staining intensity was increased in tumor cells compared with normal epithelium (40) (**Figure 2**). Finally, we investigated the expression of CEA, which has been reported to have positive staining in PDAC (41). We observed positive staining in the tumor group, as well as the PDX and organoid models (**Figure S2**). In summary, we observed a strong concordance of expression between the primary tumor, organoid and PDX models for all the protein biomarkers examined.

### DNA and RNA sequencing demonstrates molecular profiles common to primary tumor, PDX and organoid models

We next investigated if PDX and organoid models preserve the genomic and transcriptomic characteristics of their corresponding tumors. We extracted DNA from the primary tumor, peripheral blood, PDX, organoids and 2D cell lines for seven of the patients in our study and performed targeted sequencing of a 1,213 genes panel comprised of known cancer-related genes (20). We achieved an average coverage of 930X for tier 1 genes and 491X for tier 2 genes (20). Single organoids (5-9) were sequenced from each patient in order to assess the extent of intra-patient model heterogeneity. Consistent with previous reports (42-44), primary tumors from all patients had mutations in *KRAS* and *TP53* which were reproduced in PDX, organoid and 2D cell lines. Furthermore, mutations specific to individual patients such as *CDKN2A, NALCN, ZBTB16* and *PARP1* were also present in corresponding tumor models (**Figure 3A**). Remarkably, these same patterns were observed in primary tumor-derived organoids (**Figure 3B**). Genetic mutations were stable over time and passage number in PDX-derived organoids, as evidenced by sequencing (patient 2) at passages 4 and 10 (approximately 8 weeks apart) (**Figure 3C**). Interestingly, the variant allele frequency for mutations in the tumor models was higher than that of the primary tumor (median value: 57.69 and 12.44, respectively).

**Figure 3.**
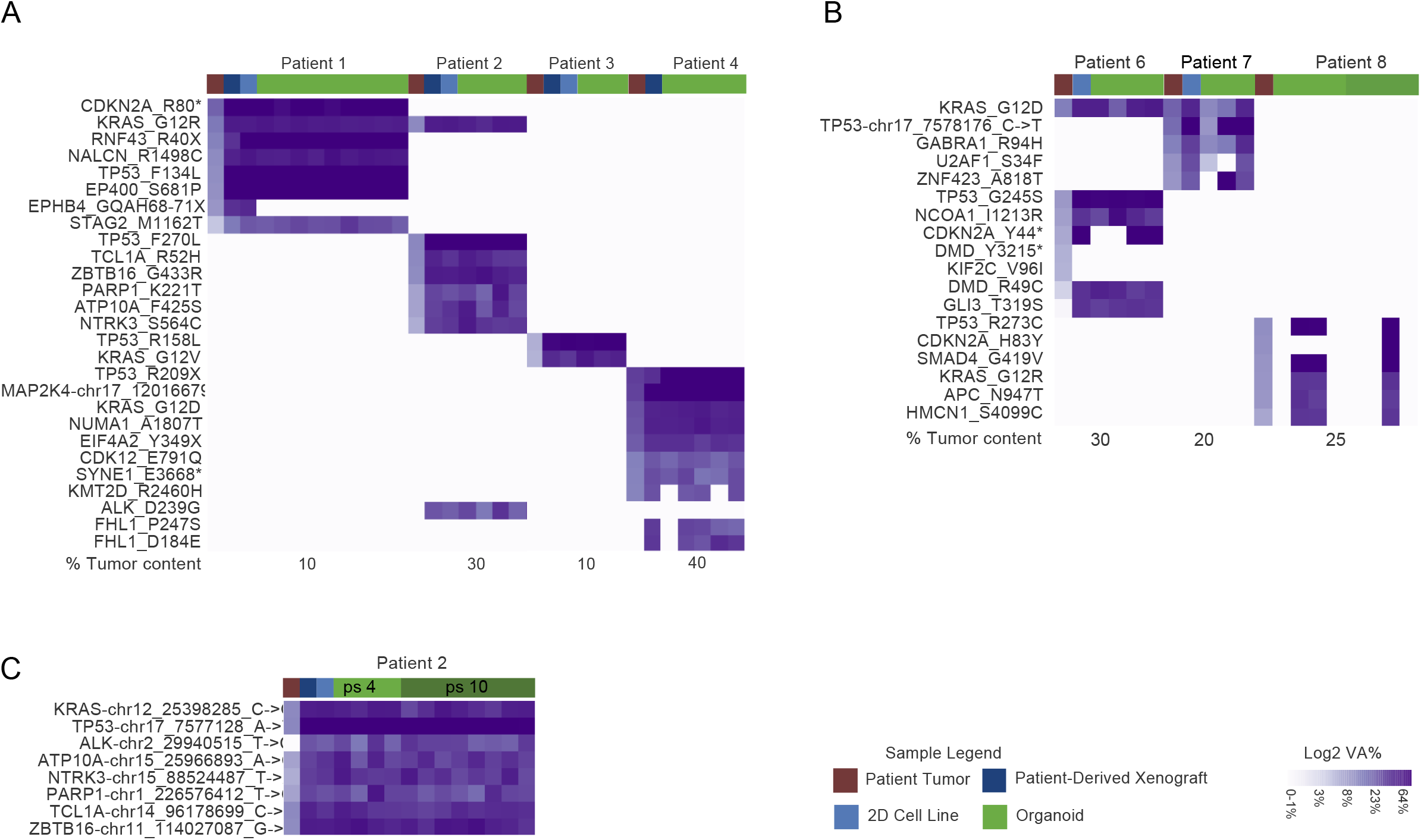
Patient specific DNA mutations are maintained by 2D cell lines, PDX and PDX- and tumor-derived organoid models. (A) DNA sequencing comparison of primary tumor (dark red), PDX (dark blue), 2D cell lines (light blue) and PDX-derived organoids (green), patient 1-4. (B) DNA sequencing comparison of the patient-derived organoid group, patients 6-8. Mutations can be detected in organoids with low tumor cell counts (tumor count 10-40%) (A and B, bottom). (C) Organoid sequences profile are shown over several passages (passage 4 vs. passage 10).

To understand differences in transcriptional profiles between PDAC from individual patients and their corresponding PDX and organoid models, we performed mRNA sequencing (RNA-seq) on the primary tumor, adjacent normal tissue, PDX models and 2D cell lines from patients 1-4. Principal Component Analysis (PCA) of the RNA-seq data indicated that the primary tumor and related models (PDX and 2D cell lines) are more similar in their gene expression profiles than to adjacent normal tissue (**Figure S3**). Results showed a high correlation between primary tumor and PDX (R squared: 0.82-0.93) and 2D cell lines (R squared: 0.51-0.90), in contrast to primary tumor and matched normal tissue (R squared: 0.09-0.6) (**Figure 4A**). To verify that the normal pancreatic tissue collected is representative of normal pancreas, we compared transcriptional profiles of normal pancreas to brain, muscle, and pancreatic tissue from the Genotype-Tissue Expression project (GTEx). These analyses verified that the normal pancreas samples collected in this study were indeed representative of larger normal pancreatic tissue cohorts (**Figure S4**).

**Figure 4.**
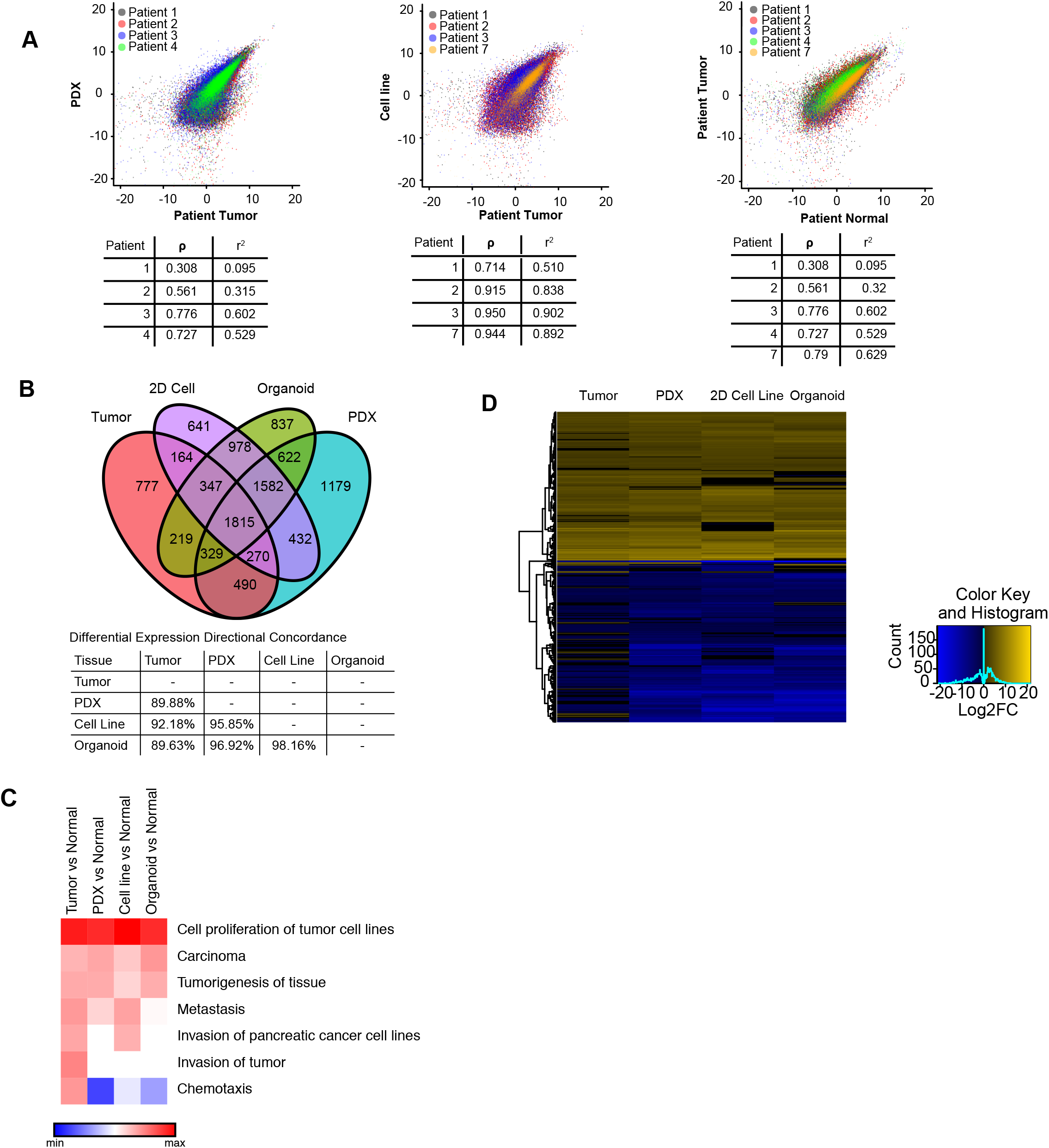
Global gene expression pattern of the primary tumor is conserved between models. (**A**) Correlation between RNA-sequencing measurements (in log2 TPM) from tumor and PDX (left), tumor and 2D cell line (middle), tumor and normal (right) for patients 1-4 and 7 are shown. Pearson correlation coefficient (p) and R square (r2) are listed (tables). (**B**) Venn diagram of all differentially expressed genes from Patient 1 comparing tumor, 2D cell lines, organoids and PDX (2-fold up or down from pooled normal, adjusted p<0.05). (**D**) Heat map of the top 200 significant differentially expressed genes shows up or down regulation in the tumor as well as the models (adjusted p<0.05). (**E**) Functional analysis of differentially expressed genes (2-fold up or down from pooled normal, adjusted p<0.05) shows signaling pathways that are up- or down-regulated across all samples (adjusted p<0.05).

To further investigate RNA expression profiles, pathway analysis was performed on the tumor, PDX, and 2D cell line RNA-seq data. We discovered that cancer-related gene expression pathways, including p53 signaling and cell cycle regulation are similarly up or down-regulated across all sample types (**Figure S5A**). Notably, upregulated p53 signaling is consistent with an increase in observed TP53 protein levels (**Figure 2C**). Differences in immune signaling dependent on model type were also apparent (**Figure S5B**), which is not surprising because PDXs are grown in immunodeficient mice and the 2D cell lines lack any immune component.

Lastly, we compared organoid RNA-seq analysis from organoids (patient 1) to the models and primary tumor. We observed a strong correlation between organoids and primary tumor (R squared: 0.66), and with both PDX (R squared 0.84) and 2D cell lines (R squared: 0.85) (**Figure S6**). The Venn diagram representation comparing tumor, 2D cell line, organoids and PDX shows that the majority of the differentially expressed genes (1815 genes) overlap between the three models and the tumor sample (**Figure 4B**). Furthermore, a majority of differentially-expressed genes are similarly up- or down-regulated across all models in comparison to the primary tumor (**Figure 4D**). Functional analysis of differentially expressed genes indicated that proliferation and cancer-related genes are upregulated across primary tumor and models when compared to normal, whereas some models show decreased expression of invasion-related genes (**Figure 4C**). Overall, we observed that 2D and 3D models maintain much of the gene expression that contributes to tumor phenotypes, although differences are apparent.

### Single cell RNA-seq reveals transcriptionally distinct subpopulations within organoids

To understand the cellular composition of the organoid models, we performed high throughput single cell RNA-seq using the Drop-seq platform (24). We sequenced the transcriptomes of 7,675 single cells derived from organoids (patient 1). We detected a median of 417 genes per cell (mean 715 genes per cell) at an average sequencing depth of ~20,000 reads per cell (**Figure 5A**).

**Figure 5.**
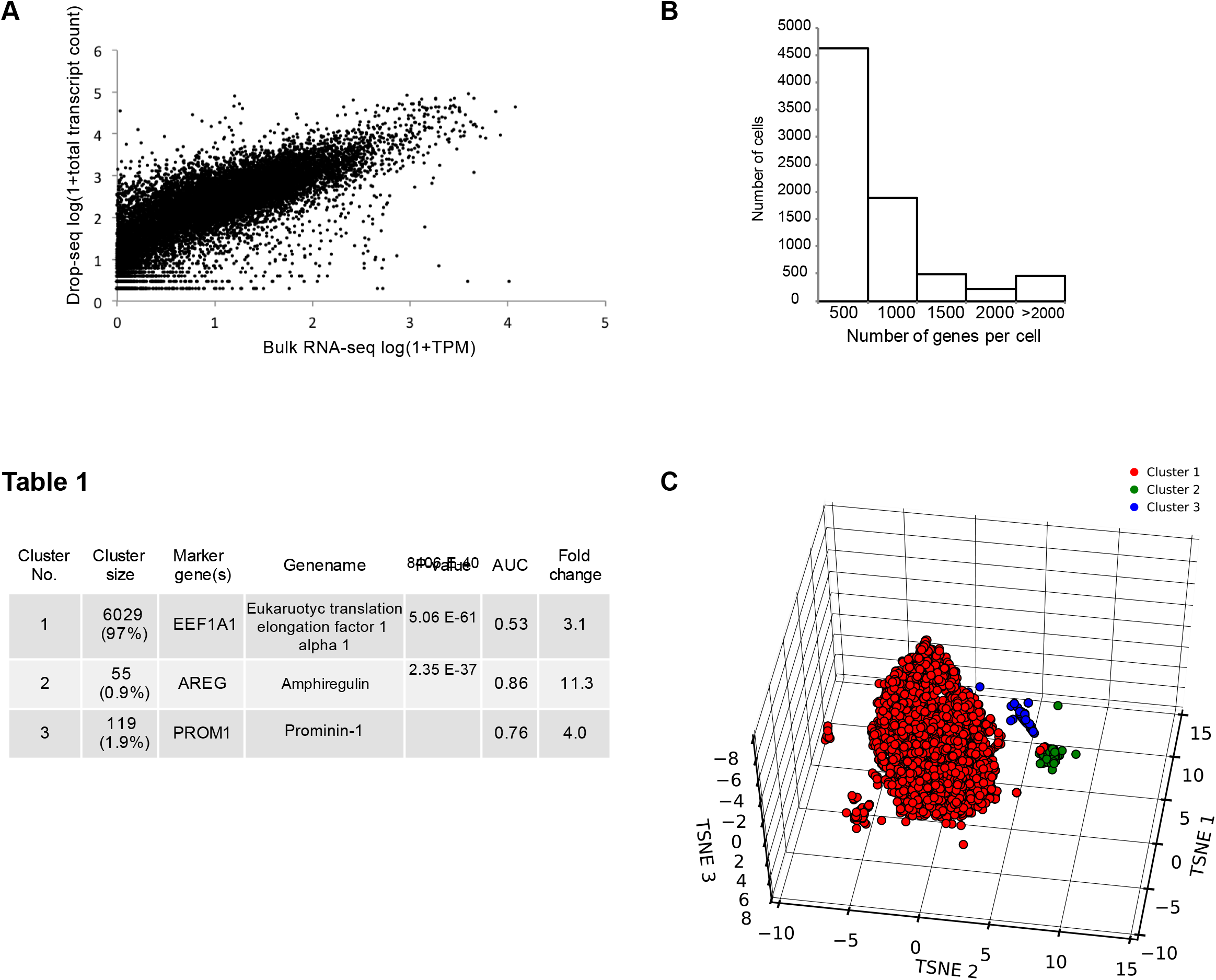
Single-cell RNA seq reveals organoids contain transcriptionally distinct subpopulations. (**A**) Correlation between gene expression measured by single-cell and bulk RNA-seq of organoids (Patient 1). The scatter plot shows the comparison between bulk RNA-seq expressed as log(1+TPM) (x-axis) and Drop-seq expressed as log(1+total transcript count) (y-axis). (**B**) Histogram shows the distribution of the number of genes detected per cell in the organoid single-cell RNA-seq dataset. (**C**) Clustering of 7675 single transcriptomes into 3 populations. A three dimensional tSNE representation (right) of the clusters predicted by topic modeling. The largest cluster (red) has been downsampled by 50% for clarity. Marker gene and p-value for each cluster are indicated in the **Table 1** on the left, AUC and fold change. A list of all marker genes for each cluster is provided in the supplementary materials (**Data file**).

We first compared Drop-seq measurements to the bulk RNA-seq data discussed above, and found the data were correlated (R squared: 0.65) (**Figure 5B**). We then clustered the single cell transcriptional results to identify distinct subgroups of cells, using a computational approach based on a class of probabilistic generative topic models. Clustering of the Drop-seq data resulted in 3 transcriptionally distinct groups of cells (**Figure 5C**). The largest cluster comprised 97.2% of the total population of cells, while the remaining 2.8% of the cells were divided between cluster 2 (0.9%) and cluster 3 (1.9%). Interestingly, cluster 3 expressed Prominin-1 (CD133), a marker gene for pancreatic epithelial stem and progenitor cells and pancreatic cancer stem cells (CSCs) (**Table 1**) (45). Cluster 2 expressed markers amphiregulin (AREG) and epiregulin (EREG), which are epidermal growth factor ligands that have been implicated in tumorigenesis and poor outcomes (**Table 1**) (46) (47). A list of all marker genes for each cluster is provided in the supplementary materials (**ListS1**). These data indicate that the pancreatic cancer derived organoids are largely clonal populations derived from the primary tumor.

### Organoids demonstrate differential responses to drug treatments

To determine if PDX- and primary tumor-derived organoid models can be used for drug response assays, we tested organoids for dosage dependent and drug-specific responses and investigated if organoid drug responses could recapitulate PDX results.

We treated organoids with gemcitabine, a chemotherapeutic agent used in PDAC, and measured resulting levels of apoptosis. Specifically, PDX-derived organoids (patient 1) were treated with three different concentrations of gemcitabine (3nM, 10nM, 30nM) for 72 hours and apoptosis was measured in real-time (**Movie S1**). Treatments caused a dose-dependent apoptotic response to gemcitabine (P =0.003 for 10nM, P=0.009 for 30nM, 3nM not significant) (**Figure 6A**). To determine whether primary tumor-derived organoid samples would yield similar results, we performed the same apoptosis assay in patient 6. Again, we observed a dose-dependent response, and interestingly, there was a full response to gemcitabine at 10nM indicating that the sensitivity threshold was higher in this sample compared to patient 1 (**Figure 6B**).

**Figure 6.**
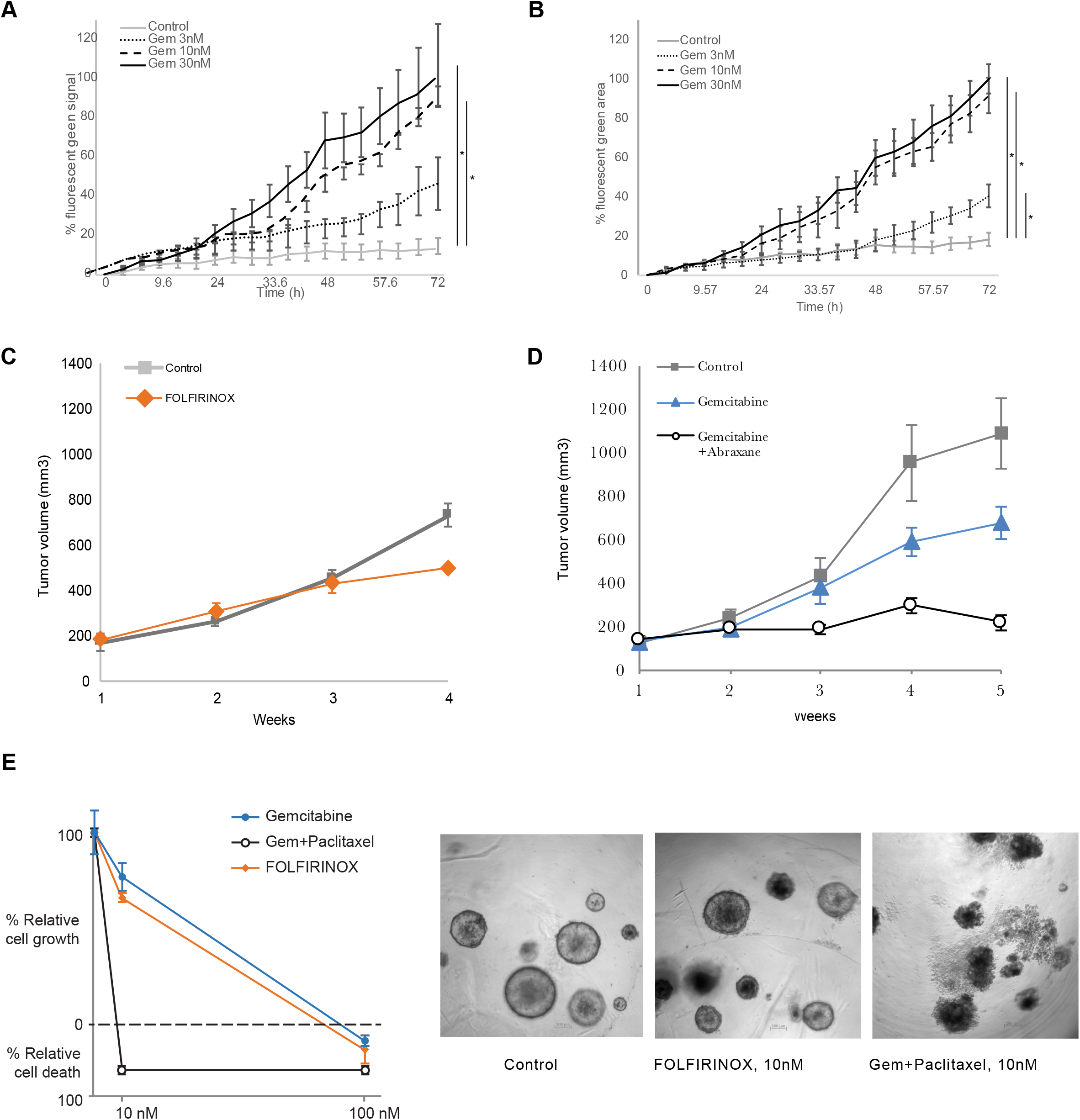
Patient-derived models respond differentially to chemotherapeutic agents. (**A** and **B**) Real-time levels of apoptosis for PDX-derived organoids from patient 1 (**A**) and tumor-derived organoids from patient 6 (**B**) were treated with gemcitabine (3nM, 10nM, 30nM). Apoptosis was measured in real-time over 72 hours (% fluorescent green signal, n=4). PDX from patient 1 (as in A), was treated in vivo with FOLFIRINOX (10nM) (**C**), or gemcitabine (10nM) with or without Abraxane (10nM) (**D**). Tumors growth (mm3) was measured over 4 weeks. Control mice received vehicle, (n=3). Points are mean tumor volume; bars, SE. (**C** and **D**). The same treatment was applied in vitro to PDX-derived organoids from patient 1. Proliferation was measured by %relatively cell growh/death (**E**) SE,*p<.001. Representative light microscopy (10x) images at 10nM are shown (right panel).

To investigate the *in vivo* drug response to FOLFIRINOX, we utilized a PDX from a patient that became resistant to FOLFIRINOX post-surgery (patient 1, **Table S1**). Following PDX-engraftment, tumor volume was measured over the course of 4 weeks. We did not find a significant benefit of FOLFORINOX treatment (**Figure 6C**). To mimic patient treatments after FOLFIRINOX resistance, we performed a second PDX experiment employing the PDX tumors that become resistant to FOLFORINOX *in vivo*. Mice were treated with a standard-of-care combination therapy for PDAC, Gemcitabine with or without Abraxane (Protein-bound Paclitaxel) (**Figure 6D**). Tumor volume was measured over 5 weeks and showed that combination therapy reduced the growth by approximately 85% compared to controls (Gem vs Con, P=0.0017), and reduced tumor growth almost twice as effectively as gemcitabine alone (Gem/Abrax vs Gem, P=0.0025). To determine if organoid models recapitulated this response, PDX-derived organoids (patient 1) were treated with the same agents followed by analysis of cell viability. A combination of Gemcitabine and Paclitaxel dramatically decreased cell viability, whereas Gemcitabine or Paclitaxel alone only reduced approximately 20% cell growth, which was not significantly different from untreated organoids (**Figure 6E**).

Overall, these data demonstrated that treatment of PDX-derived organoids recapitulates the effects of *in vivo* treatments. Therefore, organoids derived from either PDX or primary tumors are not only suitable for drug treatment studies, but results may also reflect patient-specific sensitivities to drugs.

## DISCUSSION

Here we have defined the histopathology, genetic heterogeneity and therapeutic sensitivity profiles of organoid models derived from PDAC patients. The results indicate strong concordance at the morphological and molecular levels, and are promising for the development of personalized organoid models that could be used to help guide therapeutic decisions. We have shown that organoids can recapitulate the morphological architecture of the primary tumor of origin. Specifically, we identified three major epithelial growth patterns that are found in the primary tumor and are well reflected in organoid models. These structures include simple, papillary, and cribriform morphologies that can be indicators of degree of tumor differentiation. The use of protein expression profiles in histopathological classification is used along with structural information to determine tumor type and degree of differentiation. We demonstrated that commonly used protein markers for PDAC analysis are maintained between both organoid and PDX models and primary tumors. Importantly, because organoids can be grown from a fine-needle biopsy, information such as structure and protein expression patterns may be obtainable even prior to surgical resection, opening the possibility to facilitate rapid interventions based on personalized organoid molecular, cellular, and therapeutic response characterizations.

Genetic sequencing of PDAC is challenging due to the large proportion of stromal cells in the tumor and heterogeneity of epithelial cancer cells. Since organoids have only an epithelial component, they have an amplified signal compared to primary tumor allowing deeper sequencing of tumor cells and increased detection of mutations. Previous genetic sequencing of PDAC has shown genetic similarities between the organoids with the primary tumor, although these studies have not investigated the heterogeneity of the organoid population or the stability of genetic characteristics over time. This is the first report of single organoid DNA sequencing over time using multiple passages of organoids from the same patient tumor. Our results indicate that the genetic composition of the organoids was consistent over time, for at least ten passages.

Understanding the heterogeneity of cellular states within individual patient derived organoids is a key step towards determining how they might be used as personalized models. The similarity of transcriptomes of individual cells within organoids from patient 1 are overall very similar and indicative of clonal origins, with >97% of cells occupying a single large cluster. However, we detected two small populations of cells that may offer insights into the biology of the disease. For instance, the group of cells distinguished by CD133 expression may be a stemlike subpopulation. Similarly, the group of cells distinguished by AREG expression may also be biologically relevant. Multiple studies have highlighted the role of AREG in tumorigenesis of epithelial malignancies, including pancreatic cancer, such as self-sufficiency in generating growth signals, limitless replicative potential, tissue invasion and metastasis, angiogenesis, and resistance to apoptosis. AREG is often overexpressed in PDAC and is predictive of poor outcome. Knockdown of AREG in pancreatic cancer cell lines inhibited their growth and sensitized cells to chemotherapy (48). Therefore, the sequencing of individual cells from organoids provides valuable information about clonal populations which may provide prognostic and predictive values.

Caveats for the organoid system include limitations of immune signaling and inability to test interactions with immune cells and response to immunotherapies. The development of 3D co-culture systems with an immune component is an important future step. Another weakness of the organoid system is the lack of stromal cells that exist in patient tumors, limiting our understanding of how the stroma impacts drug response. The genetic context of the stroma has been shown to be important in clinical outcome. Similar to immune cell co-culture, stromal cell co-culture is an important next step. Nonetheless, the observation of patient-specific gemcitabine responses in organoid models, as well as the differential response of both PDX and organoids to common combination therapies such as gemcitabine plus Abraxane or FOLFIRINOX, offer great promise for the use of PDAC organoids to help determine patient specific therapies. Increasing the numbers of mutational and germline genetic backgrounds represented by patient organoids, and developing methods to study immune and stroma interactions with organoids, will undoubtedly be important to fulfill the full potential of organoid models. Additionally, little is known about the differences that may be present between primary tumor-derived organoids and organoids derived from metastatic sites. Similar investigations to this study are required to determine to what extent organoids grown from metastatic PDAC tumors recapitulate the histological, proteomic, and genetic characteristics from the primary tumor and to what extent they have evolved.

In summary, our study has demonstrated that organoids are potentially valuable for drug treatment studies because they maintain distinct phenotypes and therefore respond differently to drug combinations and dosage. These models may allow for *ex vivo* drug testing of patient samples to steer treatment decisions. Furthermore, these models have potential utility for high throughput drug discovery assays.

## ACKNOWLEDGMENTS

We would like to thank Dr. Mitchell Posner and Dr. Jeffrey Matthews for allowing us to consent patients in their clinics and for providing study guidance; Dr. William Dale for providing direction in human subjects protocol development and clinical research operations; Margaret Eber, Megan Flanagan, and Teresa Barry for consenting patients and coordinating biospecimens; Marlin Amy Halder for assistance with DNA sequencing studies; Shuang Qin Zhang for consulting on therapeutic treatment of PDX models; Bradley Long and Sabah Kadri for assistance with DNA sequencing and data interpretation; and to the Friends of Jack Karp and Barbara Turf for their generous financial support of this work.

